# Polarity and mixed-mode oscillations may underlie different patterns of cellular migration

**DOI:** 10.1101/2022.10.31.514611

**Authors:** Lucie Plazen, Jalal Al Rahbani, Claire M. Brown, Anmar Khadra

## Abstract

In mesenchymal cell motility, several migration patterns have been observed, including directional, exploratory and stationary. Two key members of the Rho-family of GTPases, Rac and Rho, along with an adaptor protein called paxillin, have been particularly implicated in the formation of such migration patterns and in regulating adhesion dynamics. Together, they form a key regulatory network that involves the mutual inhibition exerted by Rac and Rho on each other and the promotion of Rac activation by phosphorylated paxillin. Although this interaction is sufficient to generating wave-pinning that underscores cellular polarization comprised of cellular front (high active Rac) and back (high active Rho), it remains unclear how they interact collectively to induce other modes of migration detected in Chinese hamster Ovary (CHO-K1) cells. We previously developed a 6D reaction-diffusion model describing the interactions of these three proteins (in their active/phosphorylated and inactive/unphosphorylated forms) along with other auxiliary proteins, to decipher their role in generating wave-pinning. In this study, we explored, through computational modeling and image analysis, how differences in timescales within this molecular network can potentially produce the migration patterns in CHO-K1 cells and how switching between them could occur. To do so, the 6D model was reduced to an excitable 4D spatiotemporal model possessing three different timescales. The model produced not only wave-pinning in the presence of diffusion, but also mixed-mode oscillations (MMOs) and relaxation oscillations (ROs). Implementing the model using the Cellular Potts Model (CPM) produced outcomes in which protrusions in cell membrane changed Rac-Rho localization, resulting in membrane oscillations and fast directionality variations similar to those seen in CHO-K1 cells. The latter was assessed by comparing the migration patterns of CHOK1 cells with CPM cells using four metrics: instantaneous cell speed, exponent of mean square-displacement (called *α*-value), directionality ratio and protrusion rate. Variations in migration patterns induced by mutating paxillin’s serine 273 residue was also captured by the model and detected by a machine classifier, revealing that this mutation alters the dynamics of the system from MMOs to ROs or nonoscillatory behaviour through variation in the concentration of an active form of an adhesion protein called p21-Activated Kinase 1 (PAK). These results thus suggest that MMOs and adhesion dynamics are the key ingredients underlying CHO-K1 cell motility.

## 1 Introduction

Mesenchymal cell migration is a spatiotemporal phenomenon that refers to one mode of cell movement characterized by the development of protrusive areas at the cell front and retractive areas at the cell rear and the requirement of energy consumption [1]. It is regulated by both extrinsic (e.g., chemokinetactic gradients) [2, 3] and intrinsic signals (e.g., Rho family of GTPases) [4, 5] that result in the spatial organization and subsequent dynamic remodelling of subcellular structures such as protein complexes termed adhesions that anchor a motile cell to its substrate and the actin-cytoskeleton [6]. It is an essential process for many physiological functions, including embryonic development [7], wound healing [8], inflammation and axonal growth during development [9]; it is also involved in pathophysiological conditions such as cancer metastasis [8, 10] and thrombosis [11]. Thus, a more clear understanding of the molecular processes that regulate cell migration will improve our understanding of a fundamental cellular physiology and also can lead to new discoveries for the treatment of disease.

Motile cells display a wide range of migratory behaviours [1]. They can bias their direction of locomotion by heading to the source of a stimulus, or randomly explore their surrounding environment [12]. Understanding the underlying mechanisms that lead to different patterns of cell migration (e.g. random, directed) is a challenging task given the large number of proteins involved. Indeed, over two hundred proteins have been implicated in regulating cell migration and adhesion dynamics with complex biological signaling pathways governing their interactions and activity [13]. Nonetheless, two members of the Rho-family of GTPases Rac1 and RhoA (referred to as Rac and Rho for the remainder of this study) have been identified as key players responsible for generating cellular polarity, comprised of a cell front and rear, leading to directional cell migration [14]. They transition from inactive (guanosine diphosphate (GDP)-bound) to active (guanosine triphosphate (GTP)-bound) forms, via Guanine nucleotide Exchange Factors (GEFs), and vice versa, via GTPase-Activating Proteins (GAPs) [4]. Activation is also regulated through their mutual inhibition that the active forms of Rac and Rho exert on each other. Together these two processes of activation and deactivation form the key signaling pathways for producing cellular polarization [4]. Active Rac is responsible for actin polymerization, causing membrane protrusion [15–18] and formation of lamellipodia (cytoskeletal projection of the membrane at the leading edge of the cell). Rho, on the other hand, is known to induce the formation of actin stress fibers and large stable focal adhesions [4, 5, 19], and is responsible for the actomyosin-driven contractions at the rear of a cell required for membrane retraction towards the nucleus [20].

It has been shown that the dynamics of Rac and Rho are modulated by an adaptor protein called paxillin [21]. This adhesion protein can be phosphorylated at its serine 273 (S273) residue by the active form of the protein p21Activated Kinase 1 (PAK) when bound to RacGTP (PAK-RacGTP) [22]. This phosphorylated form of paxillin can then bind to a protein complex formed by: the G protein-coupled receptor kinase InteracTor 1 (GIT), betaPAK-Interacting exchange factor (PIX), and PAK, and subsequently promote further Rac activation [22]. The phosphorylation of paxillin at S273 is one crucial switch for regulating fast adhesion assembly and disassembly [21–24]. Interestingly, substituting serine by alanine (S273A) or aspartic acid (S273D) generates, respectively, nonphosphorylatable and phosphomimetic mutants that directly affect not only the motility patterns of Chinese Hamster Ovary (CHO-K1) cells but also adhesion dynamics across the cell [22, 24] and paxillin co-binding with PAK in adhesion subdomains [24]. The nonphosphorylatable S273A mutant showed reduced motility, more stable adhesions and minimal co-binding with PAK compared to wildtype cells. In contrast, the phosphomimetic S273D paxillin mutant showed more enhanced motility, more dynamic adhesions and increased paxillin-PAK co-binding to adhesions [24].

The complexity of this system and how it impacts cell migration motivates the use of mathematical modelling approaches to quantitatively study this system [25]. Previous models have varied in complexity and analyzed different aspects of cell migration, but many of them focused on the Rac-Rho subsystem. In these models, the mutual inhibition seen in the Rac-Rho system was sufficient to generate the spatiotemporal pattern needed to generate cell polarity. This spatiotemporal pattern, called wave-pinning, is formed when a travelling wave in the cytosol is pinned in space, generating a concentration gradient in Rac and Rho and a polarized cell with a front and back, respectively [26, 27]. The dynamics of this spatiotemporal phenomenon was expanded by including paxillin and the GIT-PIX-PAK complex in a six-dimensional (6D) mathematical model characterizing how paxillin can affect polarity and wave-pinning [21, 23]. This work has highlighted the importance of paxillin in the small GTPases Rac and Rho activation.

Chinese hamster ovary (CHO-K1) cells have long been used to study cell motility. Despite their lack of significant polarity, CHO-K1 cells show significant protrusion and retraction leading to random migration as seen when they are placed on glass coverslips coated with extracellular matrix (ECM) proteins [28]. Persistent polarity leading to rapid directed cell migration is rare but the cells do form dynamic and stable adhesion complexes whose assembly, disassembly and stability are tightly regulated by proteins including paxillin [22]. Waves and oscillatory phenomena, in both cell tracks and membrane protrusions are frequently detected in various cell lines migrating on 2D surfaces [29, 30], so this prompted the question of whether wave-pinning alone can produce such behaviours, and if not, what other oscillatory dynamics must be involved. Much experimental evidence has repeatedly demonstrated the importance of shape oscillations in cell migration [31]. Several mathematical studies have attempted to understand how these oscillations are generated at the cell membrane level using a spatiotemporal model with multiple timescales [32, 33]. Existing mathematical models involving the Rac-Rho system typically accounted for polarized cells with a leading front and a stable back [26, 34, 35]. These models were then further expanded to couple the Rac-Rho system to the extracellular matrix (ECM), generating oscillatory dynamics that were either periodic or semi-periodic [36]. The amplitude of the oscillations in these models did not change or changed very little (e.g. in the case of semi-periodic oscillations). Therefore, implementing these models using the Cellular Potts Model (CPM), a computational discrete grid-based simulation technique that involves the modelling of the ECM as a mesh upon which simulated cells are superimposed [37], typically produced migrating cells that were either purely directional or do not migrate significantly but remain inactive with random cell membrane protrusion and retraction (depending on the amplitude of the oscillations), but not both. The fact that CHO-K1 cells do display both of these behaviours simultaneously, allowing them to explore and migrate [24, 38], suggests that other oscillatory dynamics may be at play.

In this study, we used a reduced four-dimensional (4D) spatiotemporal model that possesses three different timescales [39] to analyze motility patterns of CPM cells and how they compare to those exhibited by CHO-K1 cells. The model was both physiological, taking into account the dynamics of Rac and how it interacts with Rho and paxillin, as well as phenomenological to allow for the inclusion of two slower time scales. The model, in the absence of diffusion, produced temporal profiles that possessed both slow large amplitude oscillations and fast slow amplitude oscillations in the form of mixed-mode oscillations (MMOs) [40]. Several metrics developed in this study demonstrated that such MMOs are essential for capturing the key features of CHO-K1 cell migration.

## 2 Results

### 2.1 CHO-K1 cells display two distinct migration patterns: active and inactive

To explore the role of paxillin phosphorylation on serine 273, we studied Chinese hamster ovary (CHO-K1) cells. When we visually examined the migration patterns of CHO-K1 cells, data showed that they can be either (i) inactive (Fig. 1a), remaining mostly stationary (i.e. stalled) while attached to the substrate with very little movement in their membrane and very limited displacement away from their starting point, or (ii) active (Fig. 1b), exhibiting more migration activity by moving around on the substrate and travelling away from their starting point. When inspecting all the recorded tracks of imaged CHO-K1 cells, we found that they all exhibited one of these migration patterns (Fig. 1c), generating two independent classes of inactive and active cells displaying low or high levels of membrane activity, respectively (Fig. 1d).

**Figure 1:**
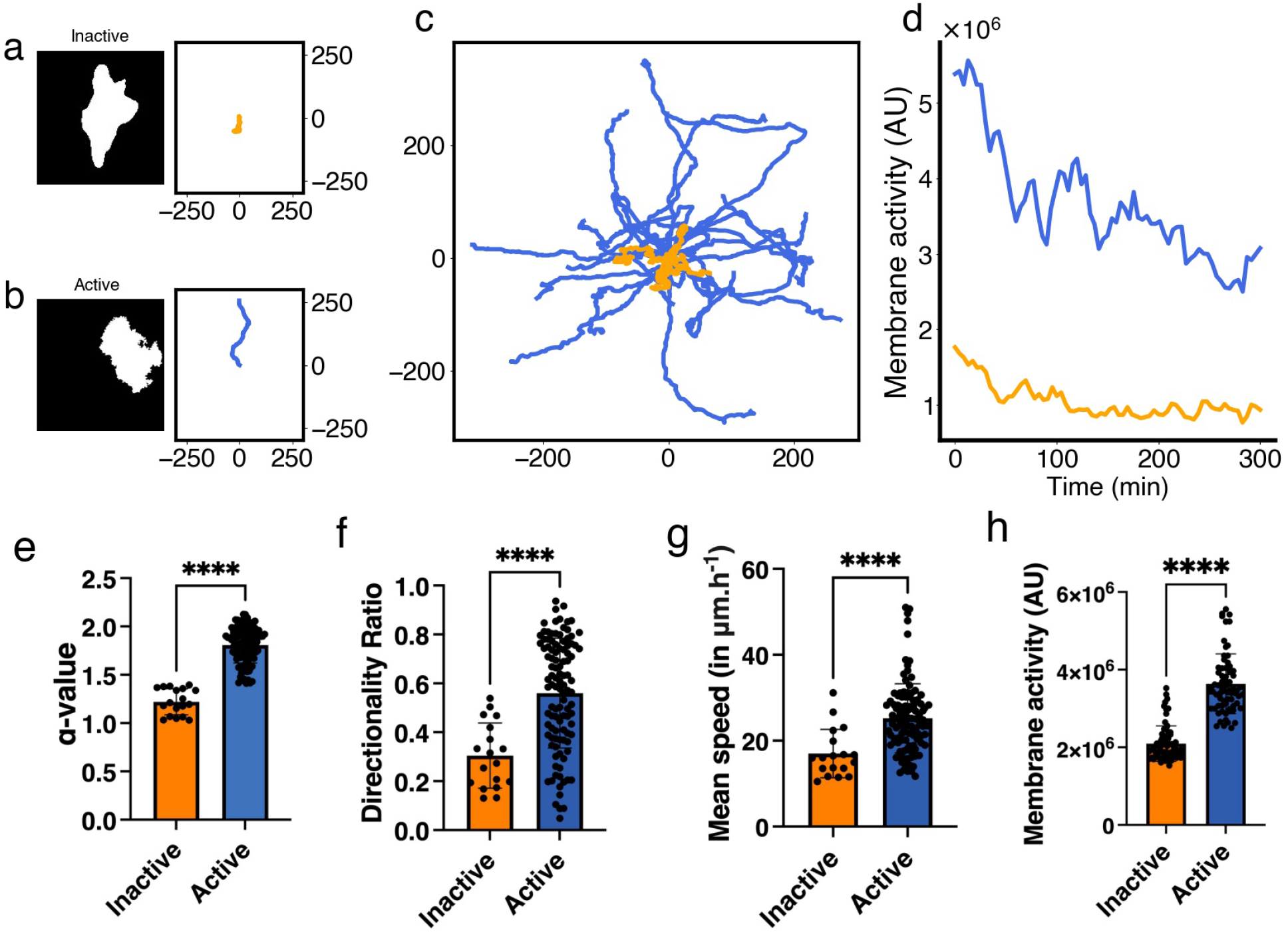
Characteristics of inactive and active classes of CHO-K1 cells exhibiting two distinct migration patterns detected using the *α*-value. (a, b) Representative binary masks (left) illustrating the shapes of CHO-K1 cells, along with their representative migration tracks (right). (c) Roseplots of a sample of CHO-K1 cell tracks (31 in total) colour-coded according to each class; orange: inactive, blue: active. (d) Time series of the level of membrane activity of two representative CHO-K1 cells, colour-coded according to each class. (e) Mean *α*-value of CHO-K1 cells in each class. (f) Directionality ratio (DR) for the entire trajectory of CHO-K1 cells in each class. (g) Mean instantaneous speed of CHO-K1 cells in each class. (h) Multiple measurements of the level of membrane activity, in term of pixel counts, of CHO-K1 cells in each class. Error bars indicate SEM. Black dots represent individual cells. Tracking data from [38] were used to produce panels c, e, f and g. Tracking data and cell images acquired for this study were used to produce panels a, b, d and h. There were n = 18 inactive cells and n = 108 active cells in panels e, f and g, whereas there were n=3 inactive and n=4 active cells in panel h.

The exponent of mean square-displacement, called the *α*-value expressed in Equation 4 (see Methods), provides a metric to automate classification of CHO-K1 cells as inactive or active. Thus, to systematically distinguish between the two classes of CHO-K1 cells detected, we first used the *α*-value to split them into two separate classes. Those that attained *α*-values below 1.4 were considered inactive, while those that attained *α*-values above 1.4 were considered active (Fig. 1e). Based on extensive exploration of the data, it was determined that a 1.4-threshold generated the best separation between these two classes. We then used three additional metrics, directionality ratio (DR), instantaneous speed and level of membrane activity to further characterize the behaviour of the two classes identified. Results revealed that the inactive class, on average, exhibited significantly lower DR values, instantaneous speed and membrane activity than active cells. Active cells displayed more wavy patterns on the membrane and oscillated in an irregular fashion (compare the blue trace that exhibits slow oscillations to the yellow trace that does not in Fig. 1d).

The differences between inactive and active cells identified here are consistent with what was previously found in the literature [28]. Using the *α*-value, we were able to demonstrate here that these two classes of cells are indeed distinct in a systematic and automated fashion. This, however, raises the question of what the underlying causes of such differences are between the two dynamic classes of cells?

### 2.1 Mixed-mode oscillations in scaled Rac concentration

It has been previously shown that wave-pinning, a spatiotemporal phenomenon that describes the propagation of a front of higher protein concentration that eventually gets pinned in space, underlies cellular polarization in motile cells [26]. It is an inherent property of some reaction-diffusion systems capable of attaining two different states for the same set of parameters but starting from different initial conditions (also called ”bistability”). Such spatiotemporal phenomenon, however, typically produces migrating cells that are persistently directional, a feature that appears not to be very consistent with CHO-K1 cell migration patterns (Fig. 1), including local Rac activities in several locations in the cytosol and wavy patterns travelling across the membrane.

To explore if other dynamic behaviours, aside from bistability, can produce migration patterns that are in agreement with those seen in CHO-K1 cells, we designed a simplified mechanistic model involving active and inactive forms of Rac (Fig 2) that possesses multiple time scales ranging from fast to slow (see Methods) [39]. In this model frame, described by the 4D model of Eqs. (3), active Rac exerts positive (negative) feedback on paxillin phosphorylation rate on a fast (slow) timescale (Fig. 2a), as well as exerts indirect positive auto-feedback (via Rho) on itself through RacGEF. The combination of the timescales in this model allows scaled concentration of Rac to display one type of oscillations that are called mixed-mode oscillations (MMOs) (Fig. 2b) [39]. In MMOs, fast small amplitude oscillations are combined with slow large amplitude oscillations that can potentially allow for local Rac activities in several locations in the cytosol to form and wavy patterns travelling across the membrane to occur.

**Figure 2:**
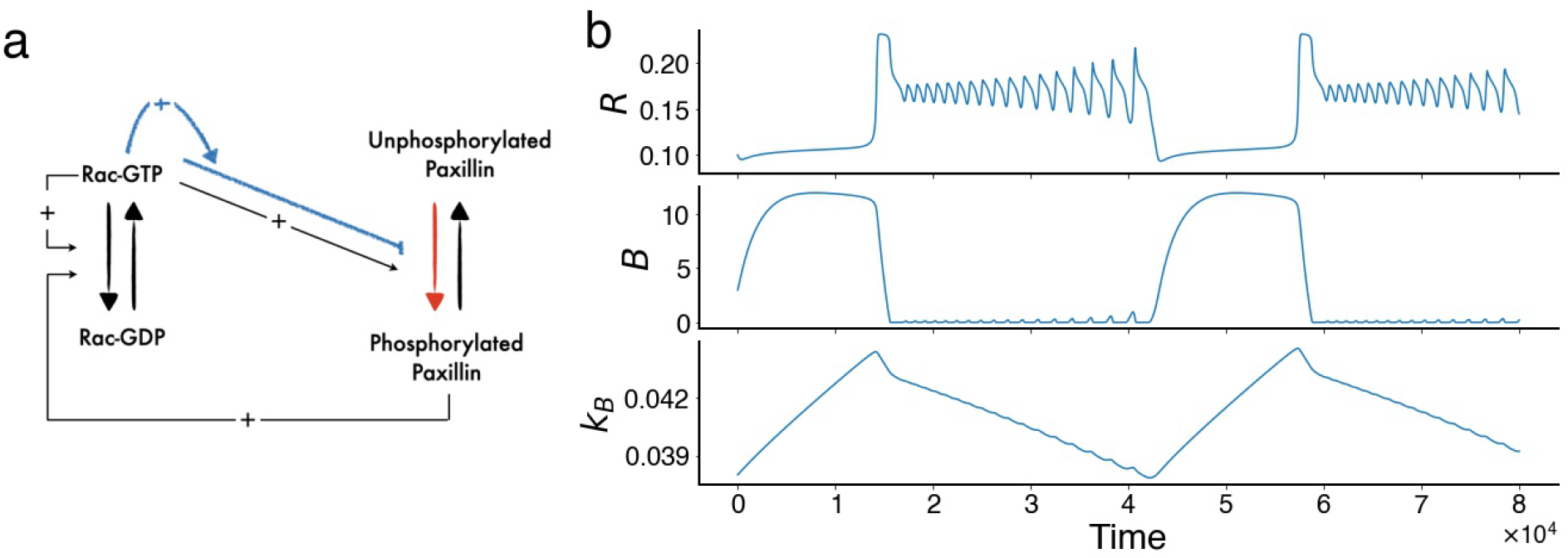
Model framework and model simulations. (a) Schematic of the semi-phenomenological four-dimensional (4D) model of Eqs. (3). Rac cycles between inactive (GDP-bound) and active (GTP-bound) forms. Rac-GTP (denoted by *R*) exerts an indirect positive auto-feedback (via Rho) on itself through RacGEF. It also simultaneously exerts positive and negative (blue pathway) feedback on paxillin phosphorylation at a fast and a slow timescale, respectively. Phosphorylated paxillin, in turn, indirectly upregulates Rac activation through RacGEF. Blue pathway: refinement of the original model [21]; red pathway: the reaction directly influenced by the half-maximum phosphorylation of paxillin (*L*_*K*_). (b) Time courses of active Rac (*R*), phosphorylation rate (*B*) and recovery rate (*k*_*B*_) in the absence of diffusion, showing the presence of mixed-model oscillations (MMOs) in *R*.

### 2.3 Cellular Potts Model (CPM) simulated cells exhibit three distinct migration patterns: directed, oscillatory and inactive

Having detected two classes of CHO-K1 cells from experimental data, we went on to explore if Cellular Potts Model (CPM) simulated cells, governed by the 4D model of Eqs. (3), can also exhibit the two distinct migration patterns. Making this link would allow us to test the hypothesis of whether the dynamic feature of MMOs, that combine slow large amplitude with fast small amplitude oscillations, underlies active CHO-K1 cell migration.

To do this, we first simulated 60 CPM cells; this produced noisy tracks with considerable variability due to the stochastic nature of the CPM simulations. When visually examining these tracks, three distinct motility patterns were discernible, one of which was inactive (Fig. 3a), exhibiting migration patterns very similar to inactive CHO-K1 cells, while the other two were active (Fig. 3b, c). The active CPM cells were either highly directed (Fig. 3c) or exhibited migration pattern that were more like patterns seen with active CHO-K1 cells (Fig. 3b). The dynamics of this latter group is governed by MMOs (and hence labelled ”oscillating” CPM cells hereafter), while directed cells are purely governed by wave-pinning [26, 27]. All the simulated CPM cells appeared to fall into one of these three classes of migration patterns (Fig. 3d) and their levels of membrane activity seemed to positively correlate with their ability to migrate (Fig. 3e). Interestingly, CPM cells in the oscillating class were found to also exhibit slow oscillations in their membrane activity, unlike the other two classes (Fig. 3e). Oscillating CPM cells on average show membrane activity that is 1.945 times higher than inactive CPM cells, a ratio that is similar to the 1.825 obtained for active versus inactive CHO-K1 cells.

**Figure 3:**
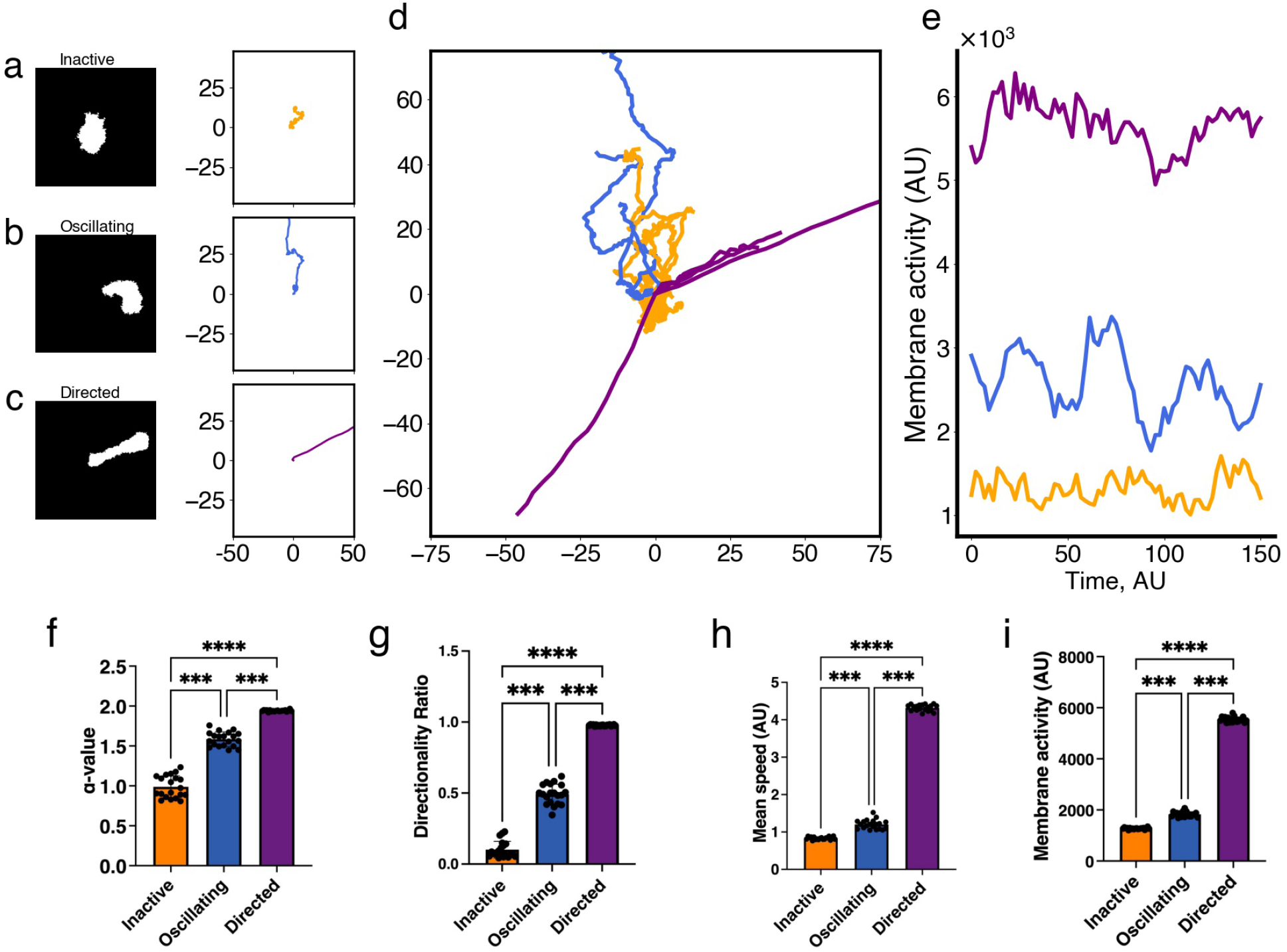
Characteristics of the three classes (n=20 for each group) of simulated cells obtained by the Cellular Potts Model (CPM) exhibiting distinct migration patterns detected by the two metrics: *α*-value, that distinguishes inactive from active CPM cells, and directionality ratio (DR), that divides active cells into oscillatory and directed cells. (a, b, c) Representative binary masks (left) illustrating the relevant shapes of CPM cells, along with their representative migration tracks (right). (d) Roseplots of a sample (n=21) of CPM cell tracks colour-coded according to each class; orange: inactive, blue: oscillatory, purple: directed. (e) Time series of the level of membrane activity of three representative CPM cells, colour-coded according to each class. (f) Mean *α*-value of CPM cells in each class. (g) Directionality ratio (DR) for the entire trajectory of CPM cells in each class. (h) Mean instantaneous speed of CPM cells in each class. Inactive cells have a non-zero mean instantaneous speed due to their inherent stochasticity, causing them to be random. (i) Mean level of membrane activity, in terms of pixel counts, of CPM cells in each class. Error bars indicate SEM. Black dots represent individual cells.

To systematically categorize the classes of CPM cells, we applied the same strategy used on CHO-K1 cells by using the 1.4-threshold in *α*-value to separate inactive CPM cells from active ones. The active CPM cells were then further divided into oscillating and directed by choosing a threshold of 0.8 in DR. This combined strategy generated a clear distinction between the three classes of CPM cells based on their *α*-value (Fig. 3f) and DR (Fig. 3g). Interestingly, the *α*-value of directed CPM cells was ≈ 2, indicating a super-diffusive motion, while the DR value of inactive cells was very low, highlighting a motion that is random (i.e. the inactive CPM cells were mostly stationary). Comparing the mean instantaneous speed (Fig. 3h) and mean level of membrane activity (Fig. 3i) between the three classes of CPM cells revealed that these two metrics positively correlated with the efficiency with which each class migrate. In other words, inactive (directed) CPM cells had the lowest (highest) mean instantaneous speed and level of membrane activity compared to other classes, an expected outcome in view of how these classes of CPM cells were defined.

### 2.4 Active Rac localization is central in defining the three classes of CPM cells

To determine what role the spatiotemporal dynamics of Rac plays in generating differences between the three classes of CPM cells, Rac activity was studied across the entire cell bodies of these simulated cells. To do this, we first considered the inactive class of CPM cells. In this class, cells always displayed homogeneous scaled concentrations of active and inactive Rac in their cytoplasm (Fig. S1a), but still attained a non-zero mean instantaneous speed (Fig. 3h). The reason for the latter is because the CPM relies on a Hamiltonian to estimate different energy configurations and tends to minimize it locally. Because the minimization includes a stochastic component, the CPM makes it possible for these inactive cells to have slight changes in membrane configurations over time, granting them the ability to move around during these energy fluctuations. Such behaviour can be seen as a random motion, with DR that drops very quickly over time to approach zero.

In the second class, oscillating CPM cells were found to exhibit episodic bursts of activity and waving patterns in their membrane (Fig. 3e). The bursts of activity were manifested as increased levels of protrusions, due to active Rac localization (Fig. S1b), that do not last for too long but are repeated periodically. Such behaviour caused the CPM cells to sometimes stretch (because of multiples protruding fronts that dynamically change localization), and to not exhibit straight migratory pattern (because of their episodic activity in protrusions and retractions) (Fig. 4a). As a result, they exhibited meandering migratory paths (Fig. 3b) with active protrusions that allowed them to migrate a lot further than inactive CPM cells (Figs. 3f and 4a, b; red arrows). The DR of their entire paths were low, but non-zero and not as low as the inactive ones (Figs. 3g). Their level of membrane activity varied depending on the phase of bursting (Fig. 4b), was positively correlated with the *α*-value (Fig. 4c) and picked up after increases in the speed of the cell; this level of membrane activity changed over time within the same track and caused the *α*-value to change meanwhile.

**Figure 4:**
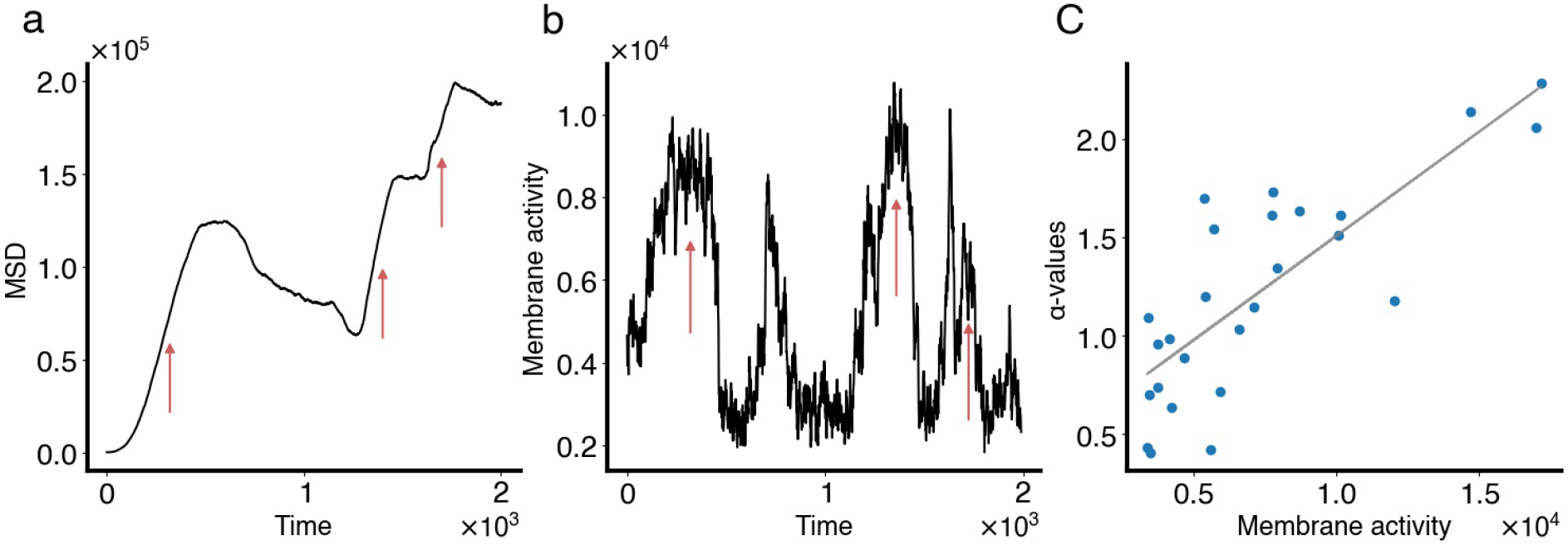
Characterizing cell movement of oscillating CPM cells. (a) MSD time series of an oscillating CPM cell. Red arrows point to periods of directed motion, when the MSD scales with *t*^2^. (b) Level of membrane activity associated with the same cell in a. Notice how periods of directed motion in the MSD correspond to periods of increased level of membrane activity, called bursts of activity (also highlighted by red arrows in this panel). (c) *α*-values plotted against mean level of membrane activity (averaged over 40 time steps) for this oscillating cell, along with the linear regression line (light gray).

Finally, directed CPM cells were found to be very polarized with active Rac accumulating in the front and inactive Rac accumulating in the rear (Fig. S1c). This extensively-studied feature of cell migration, called wavepining [26, 27], was reproducible by the model and caused the CPM cells to move in a fast and directed fashion for prolonged periods of time. Their DR values were close to 1 (Fig. 3g) and level of membrane activity were high throughout the whole simulations (Fig. 3e, i). Furthermore, their MSD indicated a superdiffusive active motion, with *α*-values close to 2.

### 2.5 Event detection in oscillating CPM cells is consistent with active CHO-K1 cells

To provide further evidence that oscillating CPM cells are representative of active CHO-K1 cells, we applied an event detection framework, designed to detect events of directionality change, on both CHO-K1 experimental and CPM simulated cells to further explore how they compare to each other.

For CHO-K1 cells, we previously distinguished active cells from those that were inactive (Fig. 1), classifying 108 tracks as belonging to the first class - active - and 18 belonging to the second - inactive. The event detection was run on the 108 active cell tracks. To ensure, retrospectively, that the detected events were indeed separating periods of directed motion, the *α*-value of each period of directedness was computed, averaged and eventually compared to the 1.4 threshold; our results indicated that, on average, the *α*-value for all periods of directedness and tracks was 2.065 ± 0.473, which is larger than the 1.4 threshold, indicating a very highly directed superdiffusive motion. As an example, for a randomly selected active CHO-K1 cell (Fig. 5a), the computed *α*-values of all detected periods of directedness were 2.069, 2.262, 2.143, 2.039 and 1.350, showing that during four out of five periods, the cell was directed. More importantly, when the *α*-values were computed using a 60-minute-long rolling window (see Methods), we found that, for each CHO-K1 cell track, the *α*-value of each period of directedness, delimited by two events, was almost always higher than its mean using the rolling window. Thus, this method allowed for the automatic identification of periods of increased directionality.

**Figure 5:**
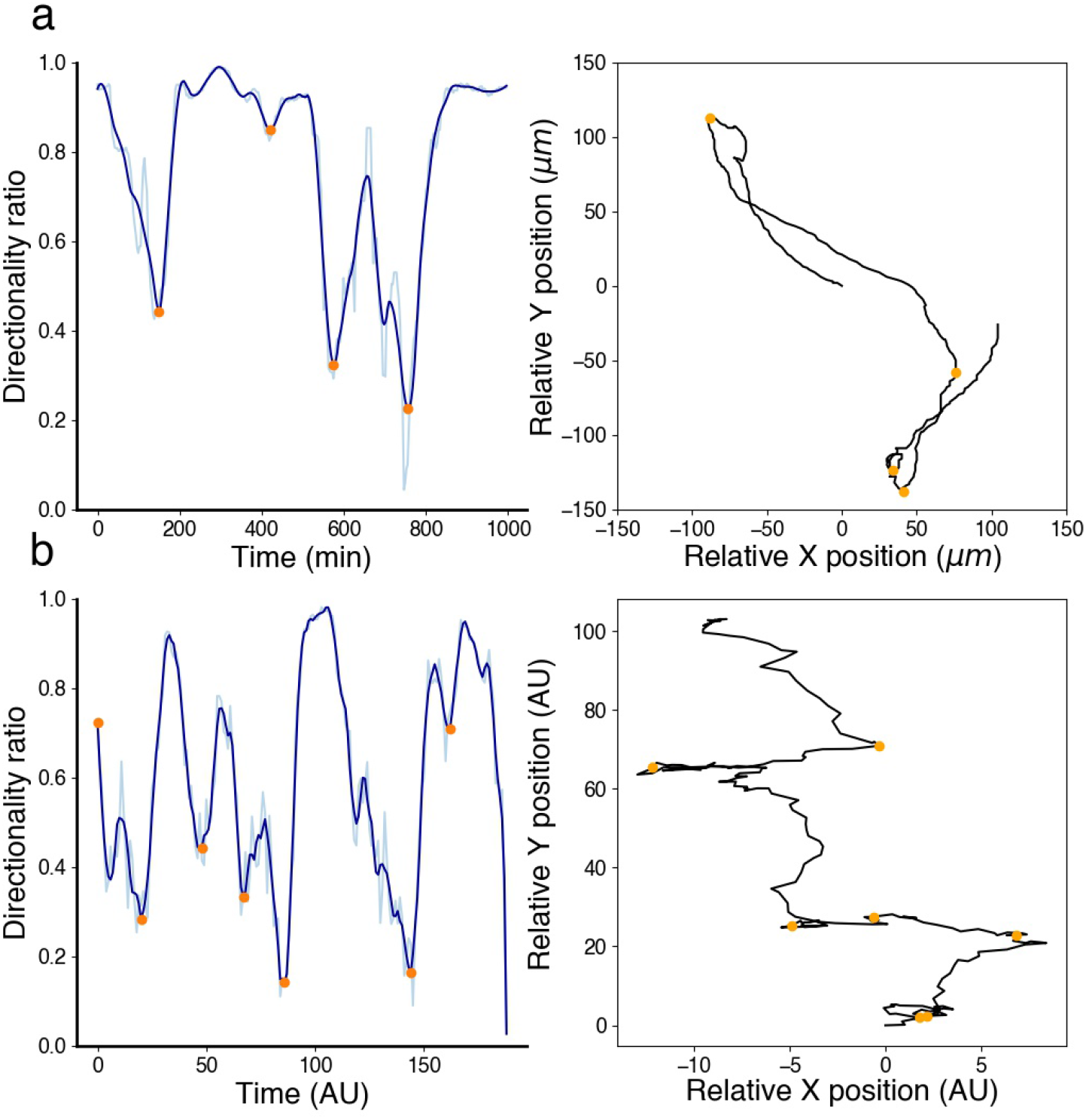
Periods of directedness and directionality change in active CHO-K1 and oscillating CPM cells. Event detection framework applied to (a) an active CHO-K1 cell, and (b) an oscillating CPM cell. Directionality ratio (left panels), computed using a rolling window of length 60 minutes (light blue) for an active CHO-K1 cell and 20 time steps for an oscillating CPM cells, allowed for event detection (orange dots). Superimposing detected events (orange dots) on cell tracks (right panel) shows a good agreement with directionality change.

The same event detection framework was then applied on 20 oscillating CPM cell tracks. Interestingly, similar dynamics concerning change of direction were observed. As in CHO-K1 cells, periods of directness were followed by clear changes in directionality. An example of such behaviour, a randomly selected oscillating CPM cell (Fig. 5b) showed that the computed *α*-values of all detected periods of directedness were 1.634, 1.56, 1.148, 2.645, 0.327, 1.724. When computing the average *α*-value of all periods of directedness and all tracks, a value of 1.851 *±* 0.80 was obtained; this value is close to that seen in CHO-K1 cells (refer to figure).

### 2.6 Mutation of Serine-273 residue on paxillin alters CHO-K1 cell migration patterns

The S273 residue can be targeted to generate phosphomimetic or nonphosphorylatable mutants of paxillin in which the serine 273 residue is replaced with aspartic acid (S273D) or alanine (S273A), respectively. Here, we investigated how such mutations alter migration patterns.

Cells expressing the paxillin-EGFP-S273A mutant showed impaired motility with typically shorter migration tracks compared to CHO-K1 cells expressing wild-type paxillin-EGFP (Fig. 6; compare panel a to b). Indeed, cells expressing the S273A mutant had significantly reduced instantaneous speed of 16.1 ± 0.658 *µ*m·h^*−*1^ and lower mean *α*-value of 1.418 ± 0.038 compared to CHO-K1 cells expressing wild-type-paxillin-EGFP whose mean instantaneous speed was 25.8 ± 0.822 *µ*m·h^*−*1^ and mean *α*-value was 1.698 ± 0.037 (Fig. 6d and e, respectively). Cells expressing the paxillin-EGFP-S273D phosphomimic, on the other hand, displayed - on average - a more active migratory phenotype than cells expressing wild-type-paxillin-EGFP (Fig. 6; compare panel a to c) with a higher mean instantaneous speed of 35.2 ± 1.93 *µ*m·h^*−*1^ (Fig. 6d) and elevated, but statistically insignificant, mean *α*-value of 1.672 ± 0.028 (Fig. 6e). These results are in agreement with previously published data [22, 24].

**Figure 6:**
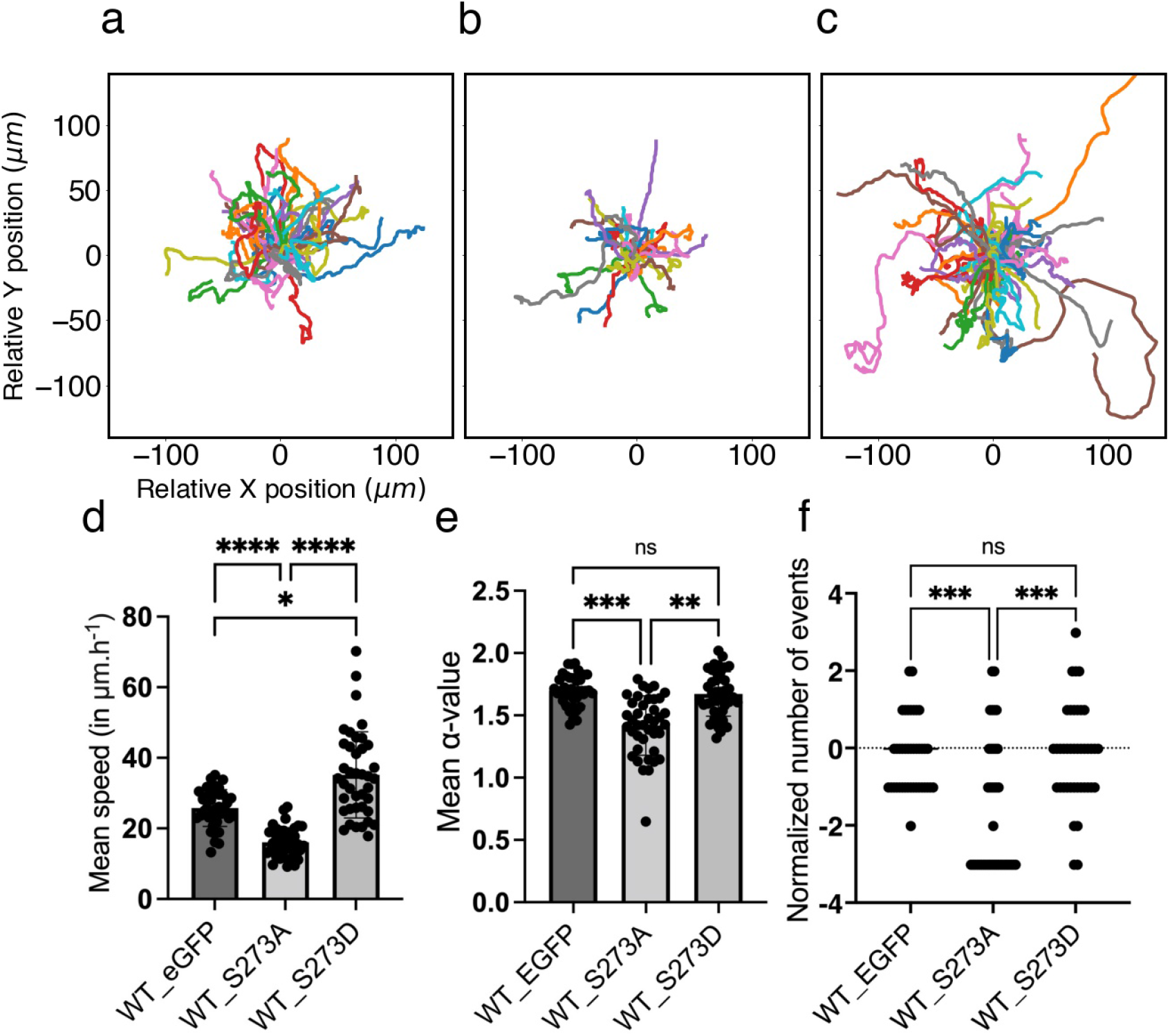
Serine-273 (S273) residue on paxillin is key to CHO-K1 cell migration. Roseplots showing cell tracks (n=40 cells for each condition) of wild-type CHO-K1 cells that were transfected with (a) paxillin-EGFP, (b) paxillin-S273A-EGFP phosphomutant, and (c) paxillin-S273D-EGFP phosphomimic. (d) Mean instantaneous speed of the same sampled cells in each condition, measured over 360 minute-long recordings. (e) Mean *α*-value of the same sampled cells in each condition, computed using a rolling window of length 60 minutes. (f) Normalized number of events of the same sampled cells in each condition computed over their entire tracks. Error bars indicate SEM. Cell tracking data from [41] were used to produce the figure.

To further explore how the migration patterns of cells expressing wild-type versus mutant paxillin compare to each other, the number of events representing changes in directionality (as defined by Fig. 5) were measured for all cells in each condition. The results were normalized by the number of events in cells expressing wild-typepaxillin-EGFP (Fig. 6f). Our results showed that cells expressing the S273A mutant exhibited a significantly lower number of events (1.93) than wild-type expressing cells (2.55) and S273D mutant expressing cells (2.62), but the latter two groups did not show statistically significant differences.

These results thus suggest that cells expressing the S273A mutant are distinct in their migration patterns, exhibiting a far less active migratory behaviour, while the migration patterns of cells expressing the S273D phosphomimic are generally equivalent to or are slightly more dynamic than cells expressing wild-type paxillin.

### 2.7 Mutant CPM cells can mimic cells expressing paxillin phosphorylation mutants by altering the model to change the sensitivity of paxillin to phosphorylation

The total active PAK concentration (i.e., total RacGTP-bound PAK) is known to be essential for paxillin phosphorylation at the S273 residue. The half-maximal activation parameter *L*_*K*_ in Eq. (2) describes phenomenologically how sensitive paxillin phosphorylation is to active PAK (PAK-RacGTP) concentration. Thus, *L*_*K*_ is a parameter that indirectly links adhesion dynamics to cell motility and causes significant changes to Rac dynamics when it is altered. For example, increasing *L*_*K*_ above ∼ 6 caused Rac dynamics to change from MMOs to being nonoscillatory, whereas decreasing it below ∼ 5.25 caused active Rac dynamics to change from MMOs to relaxation oscillations (ROs) (Fig. 7a). To investigate how altering the value of this parameter affects migration patterns of CPM cells, we simulated CPM cells at different values of *L*_*K*_ and plotted their migration tacks (Fig. 7a). Doing so revealed that setting *L*_*K*_ ≲ 6 produced actively migrating cells, while setting *L*_*K*_ ≲ 6 produced inactive or stationary cells (Fig. 7a), demonstrating its importance in defining CPM motility.

**Figure 7:**
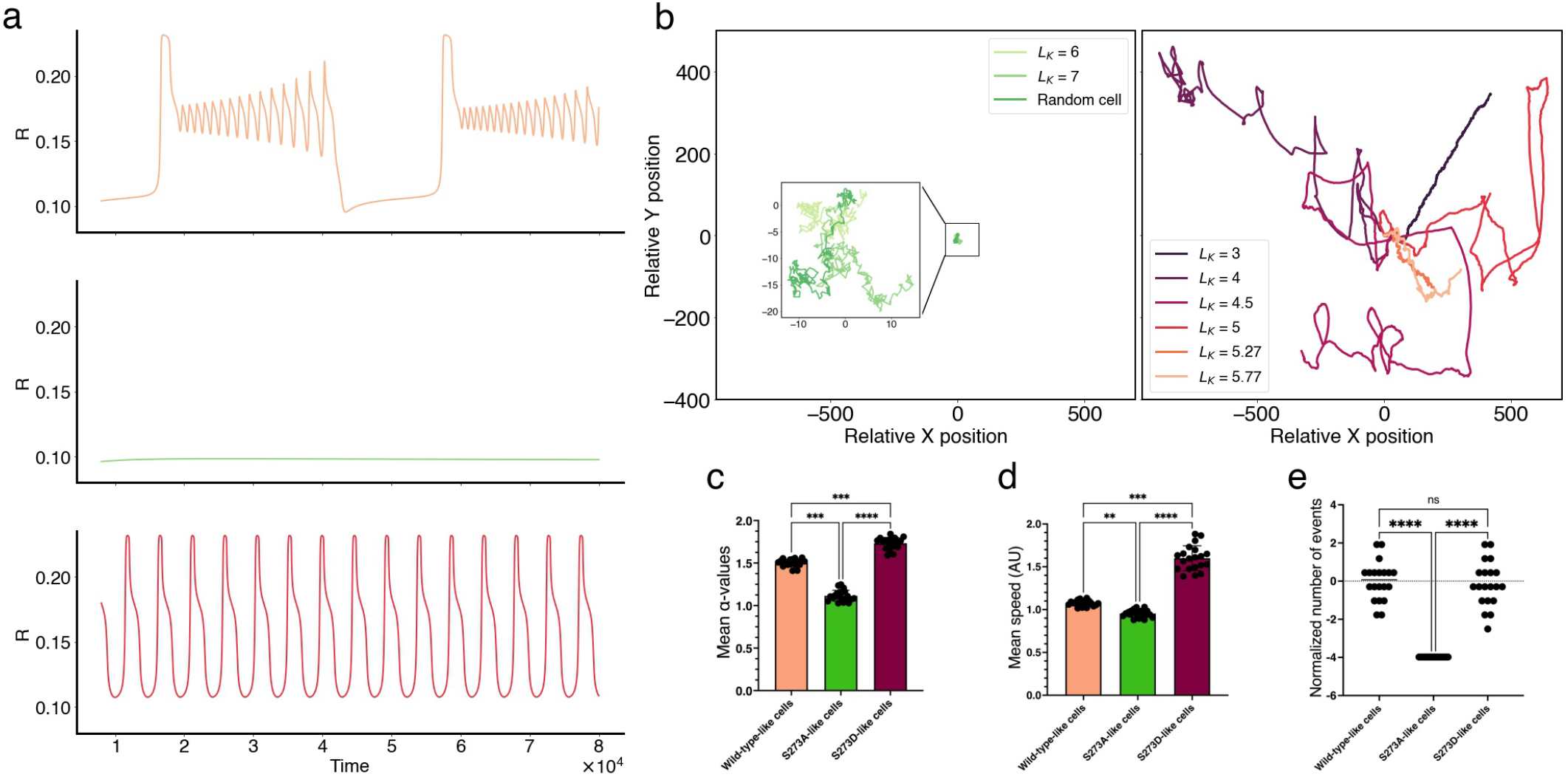
Role of PAK-dependent sensitivity parameter *L*_*K*_ in defining mutant dynamics in CPM cells. (a) Time series simulations of active Rac at default value of *L*_*K*_ = 5.77 (top) showing mixed-mode oscillations (MMOs), at *L*_*K*_ = 7 (middle) showing non-oscillatory behaviour, and at *L*_*K*_ = 4.5 (bottom) showing relaxation oscillations (ROs). (b) Migration tracks of CPM cells generated by altering only *L*_*K*_ while keeping other parameters fixed; this produced inactive (i.e., random) cells when *L*_*K*_ *>* 6 (left), or actively migrating cells when *L*_*K*_ ≲ 6 (right). The lower the value of *L*_*K*_, the further the CPM cell typically migrated. (c) Mean instantaneous speed, (d) mean *α*-values, and (e) normalized number of events evaluated over the entire simulated track from 20 cells from each of wild-type-like (*L*_*K*_ = 5.77), S273A-like (*L*_*K*_ = 7) and S273D-like (*L*_*K*_ = 4.5) CPM cells. The number of events in the S273A-like CPM cells did not vary because they all exhibited random inactive migration pattern; they were all zero and became negative when normalized. Error bars indicate SEM.

As a result, the parameter *L*_*K*_ was targeted to determine if this could generate similar migration patterns as cells expressing one of the two paxillin mutants. This was done by simulating 20 CPM cells per condition using Morpheus. In these simulations, all cells possessed identical parameter values across the three conditions except for *L*_*K*_ that was either kept at its default value of *L*_*K*_ = 5.77 to represent the wild-type paxillin expressing cells, set to *L*_*K*_ = 7 to represent the S273A phosphomutant, or set to *L*_*K*_ = 4.5 to represent the S273D phosphomimic. In the S273A-like CPM cells, the mean instantaneous speed 0.9537 ± 0.009 and alpha value 1.115 ± 0.014 were significantly lower than those for wild-type-like CPM cells (mean instantaneous speed: 1.057 ± 0.007; mean *α*-value: 1.553 ± 0.010) (Fig. 7c and d, respectively). The normalized number of events representing changes in directionality in S273A-like CPM cells (−3.981; negative values due to normalization) was also significantly lower than the value for wild-type-like CPM cells which was 0, due to normalization (Fig. 7e).

In the case of S273D-like CPM cells, however, the instantaneous speed (1.60 ± 0.032) and *α*-values (1.690 ± 0.0170) of these simulations were significantly higher than wild-type-like CPM cells (Fig. 7c and d, respectively). The former is in agreement with how experimental cells expressing the S273D mutant compare to cells expressing wild-type paxillin, but the latter is not. This seems to suggest that distinguishing between the three cell conditions using the parameter *L*_*K*_ would be better captured by using distributions of *L*_*K*_ values (centred around those used for each condition) to describe them, rather than using one specific value of *L*_*K*_ to describe all CPM cells within each condition. This is in line with cell-to-cell variation expected in experimental systems. To further validate this argument, the normalized number of events representing change in directionality was calculated (−0.1475 relative to 0 for the wild-type-like cells) and was not statistically different from wild-type-like CPM cells (Fig. 7e), in agreement with what was seen in experiments with CHO-K1 cells (Fig. 6f). This suggests that the number of events in both wild-type-like and S273D-like cells are equivalent.

### 2.8 Machine classification-based approach to validate the CPM model

Now that the CPM model captured the key features of CHO-K1 migration, the question was: could a machine classifier be trained on simulated CPM cells belonging to the three simulated conditions, then recognize and distinguish between cells expressing wild-type-paxillin, or one of the two phosphorylation mutants S273A and S273D? In other words, can a classifier be trained with the dynamic features of simulated motile CPM cells to characterize experimental CHO-K1 cells?

An artificial neural network with 3 fully-connected layers that incorporated a set of 4 metrics: mean instantaneous speed of the cell, mean alpha-value, directionality ratio and number of events representing change in directionality was generated (see Methods). The neural network, or classifier, was trained using 10 of each wildtype-like (*L*_*K*_ = 5.77), S273A-like (*L*_*K*_ = 7) and S273D-like (*L*_*K*_ = 4.5) CPM cells. Its performance was then tested on sixty (twenty of each type) randomly simulated CPM cells. The classifier demonstrated almost-perfect accuracy.

The trained classifier was then applied to track 40 CHO-K1 cells expressing wild-type-paxillin-EGFP, paxillin-S273A-EGFP mutant or paxillin-S273D-EGFP mutant (Fig. 8). In each category, the classifier identified the majority of wild-type or mutant expressing cells accurately by labelling them according to the type of paxillin they were expressing. The distribution of cell identification for each predicted condition was centred around the matching condition (Fig. 8, insets). Interestingly, a distribution shift was observed between cells expressing the S273A versus the S273D paxillin mutants. Specifically, the classifier identified most paxillin-S273D expressing cells correctly, with few recognized as wild-type-paxillin-EGFP expressing cells and very few as paxillin-S273A mutant expressing cells; the classifier did as well on the cells expressing the paxillin-S272A mutant: only one was identified as paxillin-S273D cells.

**Figure 8:**
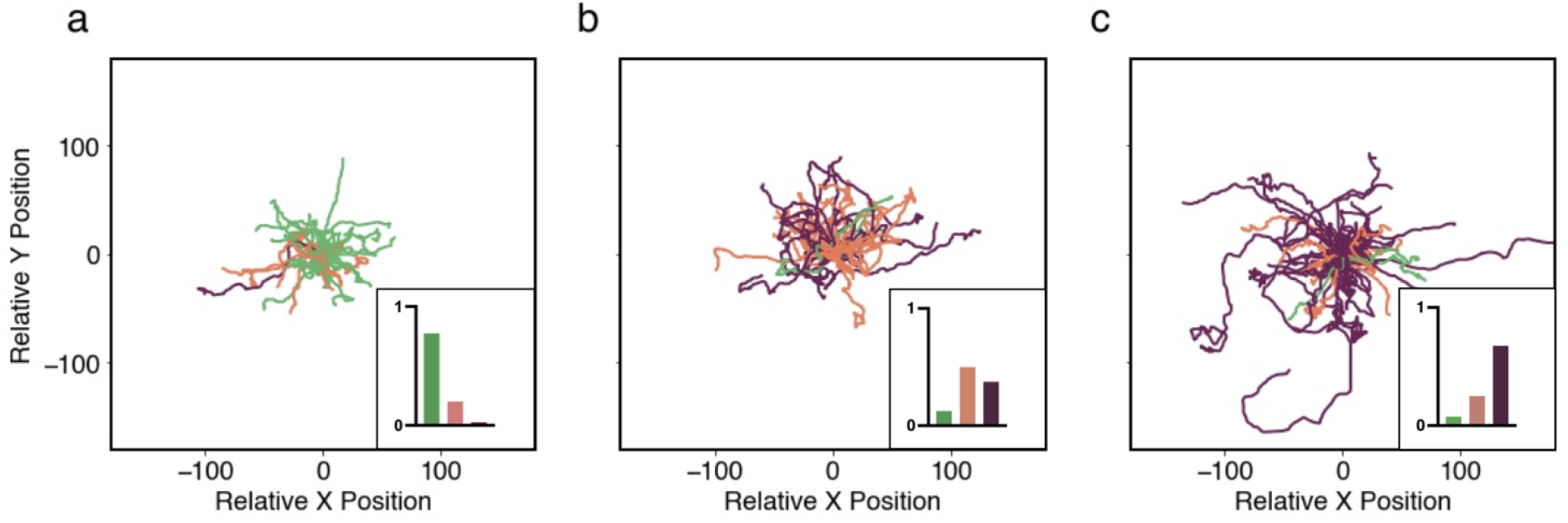
Machine classification of CHO-K1 cell tracks in three different condition. Tracks of 40 CHO-K1 cells expressing (a) paxillin-S273A-EGFP mutant, (b) wild-type-paxillin-EGFP, or (c) paxillin-S273D-EGFP mutant. Trajectories were colour-coded based on the classifier’s predictions with green corresponding to tracks classified as cells expressing the S273A mutant, orange as wild-type and purple as the S273D mutant. Insets: boxplots showing the distributions of the classifier’s predictions within each class. Tracking data from [41] were used to produce the figure.

These results thus show that the classifier is able to identify the three groups of CHO-K1 cells after training only with simulated CPM cells. This supports our hypothesis that MMOs underlying the dynamics of wild-typepaxillin-like CPM cells is key to producing expected migration patterns of CHO-K1 cells.

## 3 Discussion

The present study brought new light onto what defines cell migration in CHO-K1 cells by emphasizing the importance of the presence of multiple timescales in governing the dynamics of Rac, Rho and paxillin. The pre-existing 6D spatiotemporal Rac, Rho and paxillin model [21, 23] was simplified and modified to develop a reduced 4D spatiotemporal model to achieve this goal [39]. The model described closely the dynamics of active and inactive Rac, based on its interaction with other proteins including Rho and paxillin, but also included a phenomenological part comprised of two parameters that were turned into variables. These variables were the maximum phosphorylation rate of paxillin *B* and the recovery rate of *B* to its resting state *B*_*r*_, given by *k*_*B*_. The model incorporated, respectively, a fast and a slow positive and negative feedback from active Rac to paxillin phosphorylation (Fig. 2a).

The presence of the three timescales in the 4D model produced, in the absence of diffusion, a special kind of dynamic behaviour in the scaled concentration of Rac, namely, MMOs. MMOs combine slow large amplitude oscillations with fast small amplitude oscillations to generate interesting behaviours that we showed here to be key to CHO-K1 cell migration. By perturbing one key parameter of the 4D model, *L*_*K*_, representing how sensitive paxillin phosphorylation is to active PAK (PAK-RacGTP) concentration, it was demonstrated that the model can also produce ROs and non-oscillatory behaviour.

Having previously analyzed mathematically how MMOs are generated by the 4D model using slow-fast analysis [39], here an implementation of a Cellular Potts Model (CPM) was used to produce computer simulated cells. The spatiotemporal dynamics of these CPM cells exhibited three migratory patterns that were heavily influenced by not only the mutual inhibition of Rac and Rho, but also by the combination of slow large amplitude and fast small amplitude oscillations within MMOs. This temporal pattern resulted in CPM cells that fell into three classes with distinct modes of migration: directed with rare events of change in directionality, oscillatory with frequent directionality changes, and inactive in which CPM cells were mostly stationary.

Unlike the inactive and directed CPM cells, oscillatory CPM cells displayed meandering migration track caused by periodic bursts of activity with piece-wise directed motion interrupted by changes in directionality. The combination of slow large amplitude and fast small amplitude oscillations in MMOs were responsible for generating this behaviour. Stationary cells, on the other hand, displayed homogeneous scaled concentrations of active and inactive Rac in their cytoplasm, but still attained a non-zero mean instantaneous speed due to energy fluctuations in the Hamiltonian as seen in random motion. Finally, the motion of directed CPM cells was governed by mostly prolonged large amplitude oscillations (with minimal impact from fast small amplitude oscillations). Such migration pattern is similar to those produced by CPM cells whose underlying dynamics are governed by wave-pinning only [27] or by standard oscillatory behaviour [35, 42]. In other words, MMOs allow cells to simultaneously exhibit migratory and exploratory behaviours that standard non-MMOs or wave-pinning alone cannot, and localized active Rac activity is essential for generating the three migration patterns.

The migratory patterns of the CPM cells were then compared with the two CHO-K1 cell migration patterns detected in this study, namely, the active and inactive modes. Using a set of four metrics, including mean instantaneous speed, mean *α*-value, directionality ratio and level of membrane activity, we then characterized these two modes of migration and compared them to the ones generated by CPM cells. Our investigation revealed that the oscillatory and inactive CPM cells were consistent with the migratory patterns of active and inactive CHO-K1 cells, respectively, while directed CPM cells were not consistent with any of the migratory patterns of CHO-K1 cells. The former highlighted the importance of MMOs in CHO-K1 cell motility and suggested that wave-pinning and standard oscillations are insufficient to produce the combination of directionality change and stationarity seen in CHO-K1 cells. In addition, the results of the event detection method highlighted how similar active CHO-K1 and oscillating CPM cells are in their migration patterns, with both cells types possessing periods of directedness separated by events of change in direction. They also produced similar averaged *α*-values during periods of directedness over all tracks. This showcases that the CPM offers a good model to study and understand how CHO-K1 cell migration is regulated. CHO-K1 cells do not contain many ion channels and thus tend not to be highly polarized and do not exhibit highly directed cell tracks. Therefore, it is not surprising that the directed cell population is not observed [43].

By altering the sensitivity of paxillin phosphorylation to active PAK concentration (i.e., by perturbing the parameter *L*_*K*_), CPM cells successfully reproduced the motility patterns of CHO-K1 cells expressing one of the two paxillin-mutants: S273A and S273D. To capture the former (latter), migratory behaviour of the CPM cells were impaired (enhanced) by increasing (decreasing) *L*_*K*_. A comparison between mutant CPM and experimental CHO-K1 cells was then performed using a machine classifier trained using the simulated wild-type and mutant CPM cells. The classifier was successful in identifying CHO-K1 cells in each condition with high accuracy. This further highlighted that the CPM model governed by MMOs is sufficient to capture CHO-K1 cell migration and that perturbing adhesion dynamics reflected by the parameter *L*_*K*_ in the CPM model changes its underlying dynamics to either ROs or nonoscillatory behaviour. These latter temporal patterns are needed to capture the mutant dynamics associated with S273 amino acid phosphorylation. The classifier results also suggested that the limited number of CHO-K1 cells that were labelled differently than their actual condition may have some underlying differences in their protein network that make them different. This is not unexpected since each CHO-K1 cell expresses a different amount of the mutant paxillin protein and the cells also express endogenous paxillin that can compensate for the mutant proteins. In fact, the paxillin-S273A mutant is known to have a higher binding affinity for adhesions [24]. With this longer binding time, it is less likely to be displaced by endogenous paxillin, making the phosphomutant the dominant form of paxillin in adhesions. Thus the classifier almost perfectly identified cells expressing S273A correctly. However, the paxillin-S273D mutant has a lower binding affinity to adhesions than even the wild-type protein [24], so it can easily be displaced by endogenous paxillin. Endogenous paxillin can take on the form of the S273A or S273D mutant depending on its phosphorylation state leading to some cells being correctly identified as wild-type or S273A even though the cells are expressing the S273D phosphomimic.

The motility of S273D-like CPM cells was purely governed by ROs, allowing these cells to exhibit longer periods of directionality and to reach higher speeds. S273D-like CPM cells were very efficient in quickly changing their directionality due to the absence of small amplitude oscillations in ROs seen in MMOs. Taking a closer look at wild-type-like CPM cells, it was observed that they typically exhibited some brief random (diffusive) motions prior to directionality change, unlike S273D-like CPM cells. This random motion during change in directionality caused the discrepancy in the *α*-values that was not seen in CHO-K1 cells.

Small oscillations were visible in the time courses of the level of membrane activity in the active side of simulated CPM cells (results not shown). Limitations in the duration of imaging of CHO-K1 cells (mostly linked to the frequency at which these cells divide), however, did not allow to validate these observations on these cells. Some time courses of the level of membrane activity obtained from CHO-K1 cells showed irregular oscillations, but power spectra of these time series did not reveal anything (results not shown), because time series were too short to obtain clean power spectra. To further examine this aspect of CHO-K1 cell motility, higher time resolution imaging techniques are needed to investigate the dynamic of their protrusive activity.

Although CPMs are widely used to provide key insights into the mechanisms regulating cell motility, the nature of noise used in the simulation (white noise) can be substituted by other types of noise for purposes of enhancing the similarity with experimental biological systems. White noise does not allow for rare events or large deviations from the baseline in terms of cell perimeter or area and as a consequence, might not be closely aligned with the biological reality of cell motility. Coloured noise seems to provide a good alternative in some biological system simulations [44, 45] and its impacts could be extensively studied in the future.

The stochastic nature of the simulations can also be envisioned differently. Simulated CPM cells were very distinct across the three conditions of wild-type versus mutant paxillin and showed significant differences in all metrics, whereas CHO-K1 cells expressing wild-type paxillin and S273 mutants did not show a statistical difference in DR measurement (results not shown). This may be due to the absence of ion channels, the variability in the expression level of wild-type or mutant paxillin, the presence of endogenous paxillin or upregulation of other motility-related proteins. The classification task may be able to identify this variability and correctly identify the CHO-K1 cell behaviours. Future plans are to improve the simulation set-up by sampling the parameter *L*_*K*_ from a distribution instead of assigning each condition a single value. This would nevertheless require an estimate of the distribution of *L*_*K*_ that the parameters should be sampled from, but would produce more diverse phenotypes within a given condition.

This study involved the development of tools that can continue to be developed for simulating cell migration and varying regulation of migration through paxillin phosphorylation. The simulations accurately reproduced the known experimental outcomes. Tools were also developed for automated 1) cell segmentation 2) cell track generation, 3) identification of cell direction changes, 4) measurements of cell speed, 5) determination of type of cell movement (e.g. inactive, random, directed), 6) calculation of directionality ratios, 7) calculation of cell speed, 8) calculation of cell membrane activity and 9) classification of cell tracks.

This study demonstrates the utility of a newly developed CPM for simulated cell migration movies, provides a MMO model that can generate physiologically relevant cell migration tracks, and applies this model to further understand cell migration of CHO-K1 cells and the role of paxillin S273 phosphorylation in regulating that migration by characterizing the underlying dynamics. It uses both computational methods and mathematical modelling to achieve this goal. The approaches applied can be further expanded to analyze motility patterns in other cell types in different conditions and to include other phosphorylation pathways that regulate cell migration through the regulation of adhesions, the actin-cytoskeleton and Rho-GTPases.

## 4 Methods

### 4.1 CHO-K1 cell growth and imaging

Wild-type Chinese Hamster Ovary-K1 (CHO-K1) cells were obtained from the American Type Culture Collection (Cat. no.: CCL-61, ATCC). CHO-K1 cells stably expressing WT-paxillin–EGFP were obtained from the lab of Dr Rick Horowitz (University of Virginia, Charlottesville, VA). Stable cell lines expressing paxillin-S273A-EGFP or paxillin-S273D-EGFP were based on ATCC purchased CHO-K1 cells. CHO-K1 cells stably expressing G. gallus (chicken) paxillin-EGFP were seeded onto fibronectin coated ibidi 8-well u-slide (Ibidi, Cat. no. 80826) at a density of 2000 cells/*cm*^2^ and allowed to attach overnight. Before plating cells, the dishes were coated with a freshly made 2 ug/mL solution of fibronectin diluted in phosphate buffered saline (PBS) from a stock of 0.1% human plasma fibronectin (Sigma Aldrich, Cat. no. F-0895). The fibronectin coating was applied overnight at 4^*o*^*C*. Dishes were then washed 3x with Phosphate Buffered Saline (PBS). The CHO-K1 cells were grown in low glucose (1.0 g/L) Dulbecco’s modified Eagle’s medium (DMEM; Thermo Fisher Scientific, Cat. no. 11,885-084) supplemented with 10% fetal bovine serum (FBS; Thermo Fisher Scientific, Cat. no. 10082-147,), 1% non-essential amino acids (Thermo Fisher Scientific, Cat. no. 11140-050,), 25 mM 4-(2-hydroxyethyl)-1-piperazineethanesulfonic acid (HEPES; Thermo Fisher Scientific, Cat. no. 15630-080,) and 1% penicillinstreptomycin (Thermo Fisher Scientific, Cat. no. 10378-016,). Cells were maintained in 0.5 mg/mL Geneticin-418 (G418; Thermo Fisher Scientific, Cat. no. 11811-031,) antibiotic selection to maintain paxillin–EGFP expression. After ensuring the cells were attached to the well surface, the samples were placed in a Chamlide TC-L-Z003 stage top environmental control incubator (Live Cell Instrument, Seoul, South Korea) to maintain the cells at 37^*o*^*C* and 5% *CO*_2_ while imaging. The cells were imaged with transmitted light and fluorescence emission of EGFP on an automated Zeiss AxioObserver microscope with a Plan-ApoChromat 20X/0.8 NA objective lens and an Axiocam 506 monochrome CCD camera (Carl Zeiss, Jena, Germany, 2752 × 2208 pixels, 4.54 *µm* pixels). Transmitted light images were acquired with the halide lamp set at 3V of power and a 50 ms camera exposure time. Fluorescence images were collected using the X-Cite 120 LED light source (Excelitas, Toronto, Ontario, Canada) and a FS10 filter cube (Carl Zeiss, Jena, Germany, Ex450-490 nm, Dichroic Filter FT510, Em515-565 nm). The light power on the sample was set to 1% excitation light intensity using 10% lamp power and a 10% neutral density filter. Fluorescence images were collected with a 5 s exposure time. For protrusion measurements, images were collected at 3-minute intervals for a total of 5 hours. The resulting image timeseries were saved as a Zeiss propriety “czi” files that was later used to track the cells using a FIJI/ImageJ Manual tracking plugin. For cell tracking experiments in Figure 6 baseline measurements of cell migration, transmitted light images were acquired every minute for a total of 3 h and tracked using FIJI/ImageJ Manual tracking plugin.

### 4.2 Paxillin mutant CHO-K1 cell tracking data

Cell tracking data from previous work in the Brown laboratory was used for this study. Readers are referred to the respective publications for the experimental details, image acquisition parameters and image analysis protocols [38, 41].

### 4.3 Framework of the simplified 4D mathematical model

We have previously extended a reaction-diffusion model, comprised of the two members of the Rho-family of GTPases, Rac and Rho [16], to include the adhesion protein paxillin [21] and its effects on Rac, Rho and adhesion dynamics, through the GIT-PIX-PAK complex. The model consisted of two main modules: the mutual inhibition exerted by Rac and Rho on each other, as well as the positive feedback exerted by active Rac on paxillin phosphorylation which, in turn, exerts positive feedback on Rac activation [21]. The model was described by a 6D system of partial differential equations (PDEs) dictating the dynamics of the two proteins Rac and Rho scaled concentrations in their active (*R, ρ*) and inactive (*R*_*i*_, *ρ*_*i*_) forms, along with the scaled concentrations of phosphorylated and unphosphorylated forms of paxillin (*P, P*_*i*_) at serine 273 (S273) residue.

We subsequently simplified this model by making the following physiologically-based assumptions: (i) The synthesis and degradation of Rac, Rho, and paxillin occur on a much slower timescale compared to their reaction kinetics, implying that their total concentrations are conserved [46–48]. (ii) Rho dynamics occur at a faster timescale compared to the rest of the model, allowing us to use quasi-steady state approximation to express the scaled concentration of Rho as a function of Rac scaled concentration. (iii) The diffusion of active chemical species are very similar [49], allowing us to approximate the diffusion coefficient of these chemical species in their active/phosphorylated forms to be the same.

Taking these three assumptions into account produced a two-dimensional (2D) PDE model, given by

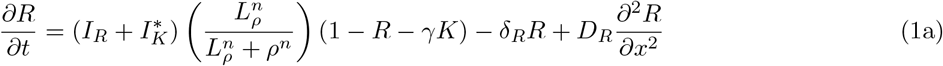

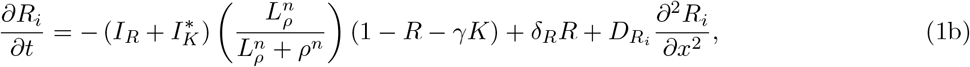

where *γ* is the ratio of total PAK to total Rac concentrations, *K* the scaled concentration of active PAK, defined by 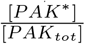, *D*_*R*_ and 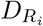 are the diffusion coefficients of *R* and *R*_*i*_, respectively, 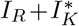 is the sum of the constant basal rate (*I*_*R*_ and the GIT-PIX-PAK complex-dependent rate 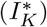) of *R* activation, *δ*_*R*_ is the inactivation rate of *R, L*_*ρ*_ is the half-maximal inhibition of *R* by *ρ* and *n* is the Hill coefficient. At steady state,

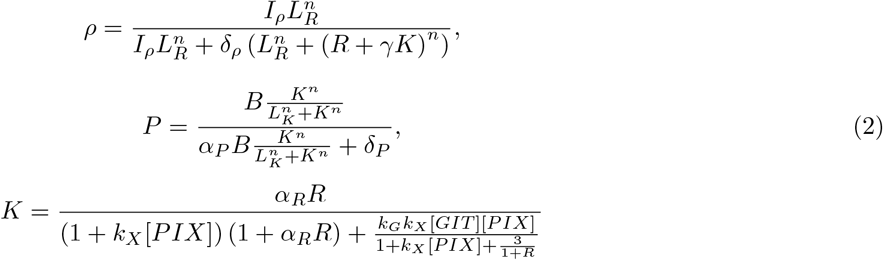

And

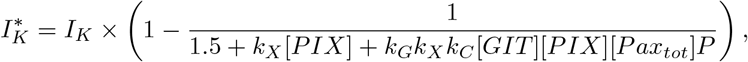

where *L*_*K*_ represents the half-maximum phosphorylation of paxillin at steady state, *I*_*K*_ the maximum Rac activation rate, *α*_*P*_ the linearization coefficient between total scaled concentration of paxillin [*Pax*_*tot*_] and scaled concentration of phosphorylated paxillin involved or not in the binding with the GIT-PIX-PAK complex, *k*_*X*_ the association constant for PIX-PAK binding, *k*_*G*_ the association constant for GIT-PIX binding, *k*_*C*_ the association constant for Pax_p_-GIT binding, [*GIT*] the concentration of GIT, [*PIX*] the concentration of PIX, *α*_*R*_ the affinity constant for PAK-RacGTP binding, *δ*_*ρ*_ the Rho inactivation rate, and *δ*_*P*_ the paxillin dephosphorylation rate [21, 23]. Technical details of how the three assumptions were implemented to simplify the model can be found in [39].

This 2D model produces dynamics that are very similar to the original 6D model [21], including bistability with respect to maximum paxillin phosphorylation rate (*B*) in the absence of diffusion, as well as the wave-pinning phenomenon, a spatiotemporal pattern with a stable front and back [26, 27]. In this study, we demonstrated that wave-pinning produces only directed motion that is not consistent with the migration patterns detected in CHO-K1 cells. To address this issue, we expanded the 2D model into a 4D semi-phenomenological model (Fig. 2a) with three timescales: fast, slow and very slow, to allow for other patterns of activity to arise, including mixed-mode oscillations (MMOs) that combine slow large amplitude with fast small amplitude oscillations within one cycle. The fast variables in this new model are *R* and *R*_*i*_, described by Eqs. (1a) and (1b), whereas the slow and the very slow variables are the two parameters - turned into variables - the maximum paxillin phosphorylation rate (*B*) and the recovery rate of *B* to its resting state *B*_*r*_ = 10 (*k*_*B*_).

The resulting 4D model is given by

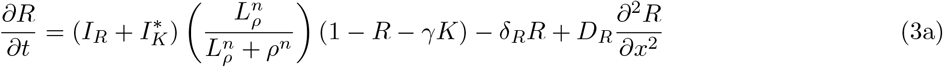

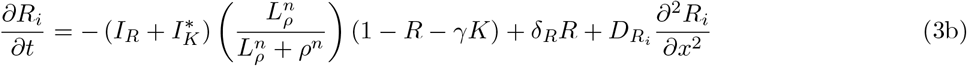

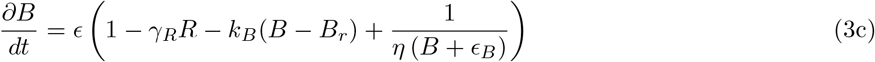

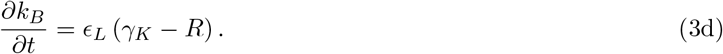

The *R* and *R*_*i*_ equations here are identical to Eqs. (1a) and (1b), respectively, where *B* was designed to vary slowly compared to *R* and *R*_*i*_, so that these fast variables can converge to their steady states before any substantial variation in *B* can occur. This slow dynamics of *B* is reflected by the parameter 0 *< ϵ ≪* 1. Mechanistically, *R* exerts positive (negative) feedback on *B* on a fast (slow) timescale (Fig. 2a). The new recovery variable *k*_*B*_, on the other hand, causes the system to exhibit oscillations of varying amplitudes and periods. It possesses the slowest timescale defined by *ϵ*_*L*_, where 0 *< ϵ*_*L*_ *≪ ϵ ≪* 1. The combination of these timescales allows *R* to display unusual dynamic properties at low values of *B*, manifested as MMOs in the absence of diffusion (Fig. 2b). We demonstrated here that such MMOs are linked to the diversity of motile behaviours observed in CHO-K1 cells. This led us to build this model in this form. Detailed analysis of MMOs produced by the model are available in [39].

Here we make four important observations: (i) Although there is a positive feedback from phosphorylated paxillin onto Rac activation, there is no direct feedback from *B* onto *R*. (ii) There is no diffusion in the last two equations, implying that they are ordinary differential equations (ODEs). (iii) It is impossible for *B* to be negative due to the fraction term in Eq. (3c) that acts as a barrier. (iv) The parameter *L*_*K*_ has a direct effect on paxillin phosphorylation (Fig. 2); it can increase or decrease the range of bistability generated by the 4D model of Eqs. (3) in the absence of diffusion when *L*_*K*_ is increased or decreased, respectively.

### 4.4 Numerical simulations of the Cellular Potts Model

The development of the Cellular Potts Model (CPM) for computer-based simulations of migrating cells has provided an excellent tool to analyze cellular motility under varying conditions. This discrete grid-based simulation technique involves the modelling of the extra-cellular matrix (ECM) as a mesh upon which simulated cells are superimposed in the form of compartmentalized discrete objects that can evolve, grow or divide on the surface of this discretized mesh. A Hamiltonian function *H*(**F**; ***λ***) is typically designed to account for forces **F**, such as pressure caused by actin filaments growth that push the membrane outward, peer contact, or external cues depending on environment-related parameters with different weights defined by ***λ***. This design is used in Monte Carlo Markov Chain (MCMC)-like simulations to mimic cellular movements, membrane deformation or even cell division, employing classically a stochastic energy minimization principle. In these simulations, the deformable area (or volume, depending on the dimensionality of the simulations) representing the cell has no pre-determined shape and movement, except for physical limitations such as elasticity limit and only responds to energy variation and physical forces.

We used *Morpheus*, a highly flexible CPM simulator software [50], to simulate a migrating cell. The software allows users to simulate cells under different experimental conditions, parameters values and dynamics, and to account for diffusive molecules in these cells. Our main motivation for choosing this particular CPM was its multi-scale component and its applicability to models expressed as reaction-diffusion equations. The software provides a tool to simulate multi-scale multicellular systems (coupling ODEs, PDEs and CPM in 2D or 3D grid), and to solve the resulting models numerically using the stochastic simulation algorithm. The system described by Eq. (3) requires this multi-scale numerical simulation method Morpheus to elucidate its underlying dynamics.

In our settings, we set the Hamiltonian function to be

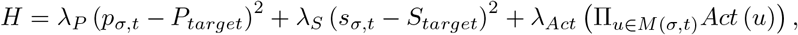

where *σ* denotes the position of the cell at time *t*, the weights *λ*_*P*_ and *λ*_*S*_ describe the deformation resistance with values tuned to allow the CPM cell to exhibit reasonable membrane elasticity properties [51, 52], *M* (*σ, t*) denotes the set of lattices that form the membrane of the cell and the *Act* function measures the amount of actin activity at a specific location. In our general CPM implementation, we substituted actin activity by its equivalent counterpart: the active Rac activity represented by *R* [53]. *All lattice sites outside the cell in this implementation were assumed to have a nonzero, but very small (0 < ϵ*_*R*_ *≪* 1) level of *R*; this makes Rac activities in these lattice sites negligible. Based on that, the software updates membrane localization by favouring membrane expansion in the regions where *R* is high, i.e. Rac activity is the highest, causing membrane protrusions. The level of *R* is each lattice site in the newly updated configuration of the cell is then dictated by the software.

### 4.5 Cell tracking

CHO-K1 cells were manually tracked in FIJI/ImageJ (NIH, Bethesda, MD) using the manual tracking plugin by clicking on the centre of the cell frame-by-frame for each motile cell in the image time series. The centre of the CHO-K1 cell nucleus was used as the reference point for each cell. Important to note that no significant difference was detected when tracking a CHO-K1 cell using its centre of nucleus or centre of cytoplasm or centre of mass. User bias was minimized by having several authors track the CHO-K1 cells. Cell tracking data from work that was previously published by the Brown laboratory were also used for these studies. Tracking data for WT-paxillin-EGFP expressing CHO-K1 cells were from [38] and CHO-K1 cells expressing EGFP paxillin fusions of WT, S273A or S273D paxillin constructs were from [41]. Simulated CPM cells were automatically tracked by Morpheus. There are slight differences across studies in image acquisition methods (i.e. transmitted light versus fluorescence imaging) and settings and cell tracking protocols. Details in the methods section refer to new experiments conducted for these studies and tracked using the methods described above. The reader is referred to the two aforementioned studies for the details of cell tracking for the appropriate figures. The *x* and *y* position data for each CHO-K1 and CPM cell track was then exported to Python. Rose plots were created by superimposing the starting position of each cell track on the origin (0, 0). The speed of each cell was then calculated by taking the mean of the distance travelled between each time point over the imaging interval, generating the metric ”instantaneous speed”. The data shown represents the mean ± standard error of the mean (SEM) for all CHO-K1 and CPM cells analyzed.

### 4.6 Comparison framework

In the absence of diffusion, the new simplified model, given by Eqs. 3, exhibits not only bistability (and thus wave-pinning in the presence of diffusion in *R* and *R*_*i*_), but also relaxation oscillations (ROs) and mixed-mode oscillations (MMOs) [39]. Thus one would expect that these diverse temporal profiles will generate different migration patterns in the simulated CPM as long as the dynamics are governed by Eqs. (3). To quantitatively verify if such migration patterns of the CPM are similar to those exhibited by motile CHO-K1 cells, an analysis framework was built to fully characterize and compare motile behaviours between simulated cell tracks and tracks from experimental timelapse movies of cells. The framework compares migration patterns by focusing on speed, persistence, directionality and membrane activity as defined below.

#### 4.6.1 Analysis of paths: metrics for quantifying cell movement

Motile cells can exhibit random motion in their migratory patterns. However, some cells may also display active migratory patterns that are not due to biological noise. Quantitative approaches can be employed to distinguish between the two. In normal diffusion, the mean-square displacement of a diffusing cell, defined by

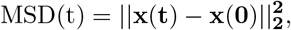

where **x**(**t**) is the position of the cell’s centre of mass at time *t*, is proportional to *t*; this applies to a cell whose motility is not driven by any concrete force but rather by the noisy fluctuations in the cell’s shape. However, diffusion can be hindered or enhanced, making MSD proportional to some power of time *t*^*α*^, given by

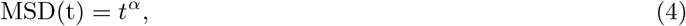

with *α <* 1 (*>* 1) indicating hindered (enhanced) diffusion.

For directed motion, MSD becomes proportional to *t*^2^ [54]. For this reason, it is more common to consider the mean-square displacement through time in log-scale and use the slope of the curve (referred to hereafter as ”*α*-value”) as a metric to determine the level of directionality [55, 56]. When *α* is close to 1, the motion is random, when *α >* 1, the motion is said to be superdiffusive, indicating that there is an active force driving directionality, and when *α <* 1, the motion is said to be subdiffusive or the cell is somehow confined. For an actively migrating cell with a speed *v*, the trajectory of this cell will simultaneously exhibit both superdiffusive and diffusive (random) motion because of the presence of biological noise. Here, the *α*-value was used to automatically classify their activity.

As with every metric, MSD has its pitfalls. Specifically, a cell that follows a curved path while maintaining an approximately constant distance from its starting point would have the same MSD as that of an immotile cell. To resolve this issue, another metric called the directionality ratio (DR) that quantifies the straightness of motion [57, 58] was also used. It is defined to be the ratio between the straight-line length from the starting point to the endpoint of the migration path, *d*, and the total length of the path along the trajectory, *D*, i.e., DR= *d/D*. According to this definition, it follows that a DR is roughly 1 for a straight trajectory and much closer to 0 for an almost stationary trajectory. Directionality ratio has its own limitations because it may attain small values for trajectories that exhibit piece-wise directed motion. In this case MSD of such trajectory should be further assessed.

#### 4.6.2 Detection of changes in direction

To characterize cell movement, a two-step approach was implemented. In the first step, an approximate value of the prevailing *α*-value over time was computed by applying log-log regression on the MSD and measuring the slope. This was done on every 20-min-long segment of the recording for CHO-K1 cells (70 segments in total), or 50 time-step-long segments for CPM cells (350 segments in total), and then averaged. A threshold value of *α* = 1.4 was found to be the most suitable in differentiating between active or motile cells (when *α >* 1.4) versus those that are inactive or moving randomly (when *α <* 1.4).

Trajectories of active CHO-K1 cells are typically ”piece-wise directional”, i.e, they exhibit a sequence of directed motions (referred to hereafter as ”periods of directedness”) separated by events at which cells abruptly change directionality. To ensure an unbiased and self-consistent detection of all these possible changes in direction or ”events”, we applied the second step in characterizing cell motion. In this step, a 60-min rolling window for CHO-K1 cells and 50 time-steps for CPM cells was used to compute the DR of a portion of the trajectory; the rolling window was then shifted by 1-min (one time-step) in the former (latter) and the computation was repeated. During periods of directedness, DR values are high (essentially above 0.8). This means that when the rolling window is overlapping with an event and is no longer purely covering one period of directedness, a significant drop in DR occurs. A minimum DR is thus attained when half of the window is in the preceding period of directness, relative to the event, while the other half is in the following period. By computing the DR values and applying non-frequency-based smoothing [59] of the DR time series, pronounced dips in times series were detected. These dips represent the events of abrupt change in directionality.

Note that tracks of inactive cells also exhibit fluctuations in DR with spikes and dips that are quite frequent. This is because DR does not include any normalization in terms of travelled distance or speed. Indeed, any Wiener process would have a noisy DR time series that sometimes reaches high values. Fortunately, these inactive cells also show very low *α*-values. Such feature can be further used to distinguish between active and inactive cells, along with the computation of confidence interval for the DR. This was how we overcame the main difficulties in the event detection task.

#### 4.6.3 Cell protrusion and shape

To investigate cell protrusion and shape in CHO-K1 and CPM cells, ”binary masks” of these two cell types were extracted [60]. These masks indicate where the entire body of the cell lies, allowing for visual identification of protrusions in the membrane and distortion or stretching in the general shape of a CHO-K1 and CPM cell (Fig. 9).

**Figure 9:**
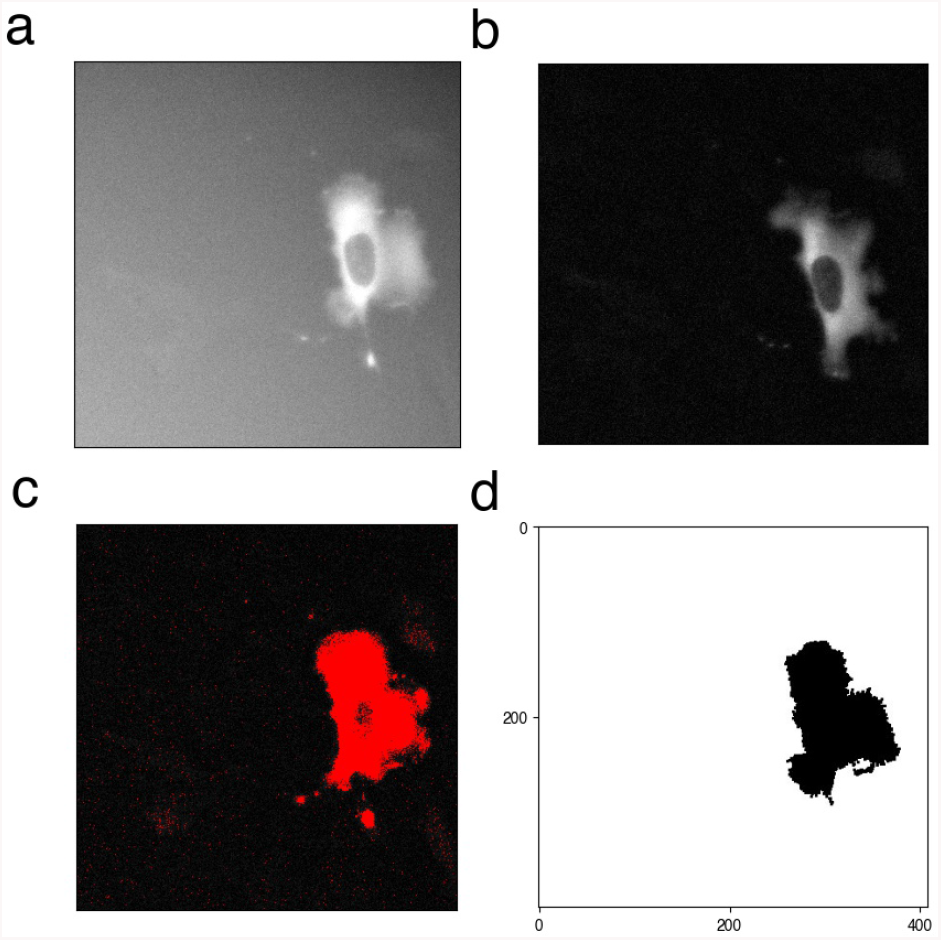
Image processing of motile cells. (a) A widefield fluorescence microscopy image of a CHO-K1 cell expressing WT-paxillin-EGFP. Such images often have a light halo and noisy background, that are (b) filtered to obtain a high signal-to-noise image, and then (c) thresholded to generate a binary image and detect the entire cell body by applying the ”find-particle” Fiji/ImageJ plug-in, resulting in (d) a binary mask of the cell. For the CPM cell, the binary masks in step d are generated directly without the need for the first three steps a-c because, by definition, the simulated images have pixels with signal and pixels that have no signal.

Fiji/ImageJ was used to generate binary masks of CHO-K1 cells by applying a classic image-segmentation framework. The cells appeared as fluorescence intensity on a dark background; pixel background intensity signal was subtracted using the ”Subtract Background” plug-in with default values; the brightness and contrast were adjusted to visualize the cells and images were thresholded to obtain black-and-white binary images using the ”Threshold” plug-in, with the default settings except that ”dark background” and ”Otsu” filter settings were selected. The ”Find -particles” algorithm was then used to identify the near-circular objects in the binary images with default settings except that ”include holes” and ”exclude on edges” were selected. The range of circularity was set to [0.3 − 1.0], minimum size to 1000 (in term of pixel count), allowing for CHO-K1 cell detection. With CPM simulations, the exact lattices (or pixels) occupied by the cell were known, making the exact detection of the cell body and generation of binary masks more straightforward. Binary masks were obtained for different motility patterns detected in CHO-K1 and CPM cells.

From the binary masks of CHO-K1 and CPM cells, changes in cell area due to protrusions and retractions in the membrane were then visually assessed. Here, we subtracted consecutive cell binary masks to count (in pixels) the size of deformations from one image to the next, in order to measure the metric ”level of membrane activity”. When a CHO-K1 or CPM cell moves forward, it is not sliding on the substrate, but rather dynamically extending its edge by forming a protrusion and retracting its rear. This leads to a displacement, and the mask subtraction is supposed to highlight these two sites of membrane deformation, i.e. the front and the rear. If masks were to be realigned, the level of membrane activity would then be close to zero, especially during periods of directedness in which CHO-K1 or CPM cells roughly preserve their shapes during migration. That is why masks were not recentred to align the two barycenters of a cell.

### 4.7 Statistical analysis

Statistical analysis was performed using PRISM software (GraphPad). Mann-Whitney tests were performed. Statistical significance was accepted at *P *<* 10^*−*1^, **P *<* 10^*−*2^, ***P *<* 10^*−*3^ and ****P *<* 10^*−*4^. Linear regression was performed using the standard method of ordinary least squares, and run on Python with its implementation from the *sklearn* Library.

### 4.8 Machine classifier

We chose to implement our classifier as a neural network, using the simulated wildtype-like, S273A-like and S273D-like CPM cells for training. As demonstrated below, the set of metrics already established in this study was quite effective in capturing the differences between the three conditions of CHO-K1 cells. We therefore chose to restrict ourselves to a very simplified architecture for the neural network; this allowed the algorithm to use the metrics efficiently and adequately with limited processing, as well as facilitated the interpretation of the results produced while reducing the risk of overfitting. The architecture consisted of 4 fully-connected layers, with a rectified linear unit (ReLU) function. The network was given the following four metrics: mean instantaneous speed of the cell, mean alpha-value, directionality ratio and number of events representing change in directionality. It must be pointed out that the units of the speed are different between the CHO-K1 cells and the CPM cells, whereas all other metrics share the same units. For this reason, we normalized the speed across all samples and kept all the other metrics unchanged. Varying the classifier, the training set and the architecture of the network, we always obtained an accuracy above 90 percent. With the adopted architecture of 3 layers, ReLU activation function and a randomized choice of 10 CPM cells per condition (wildtype-like, S273A-like and S273D-like) to form the training set, 95% was the minimum accuracy obtained, and it often reached 100 percent.

### 4.9 Software

The CPM was simulated using Morpheus (an open source software developed at the Technische Universität Dresden [50], available here). Its graphical user interface supports the entire workflow from model construction and simulation to visualization, archiving and batch processing. Plotting and classification were done using Python 3.8.12. For the machine classifier, we used the implementation of Multi-layer Perceptron classifier from the Python library Scikit learn; it was trained using the default value setting except for the choice of the solver (selected to be ‘lbfgs’), and the regularization parameter (set to be *α* = 10^*−*5^). The codes used for producing the simulations can be obtained online [61].

## Acknowledgments

We thank Alexander Kiepas for providing the CHO-K1 tracking data that were part of a previous manuscript Kiepas A, Voorand E, Mubaid F, Siegel PM, Brown CM. Optimizing live-cell fluorescence imaging conditions to minimize phototoxicity. *J Cell Sci*. (2020) doi: 10.1242/jcs.242834. PMID: 31988150 for further analysis.

We thank Abira Rajah for providing the tracking data for cells expressing the wild-type-paxillin-EGFP, paxillin-S273A-EGFP or paxillin-S273D-EGFP that were previously published in her master Thesis *The paxillin serine 273 phosphorylation signaling pathway positively regulates cell migration, persistence and protein dynamics*, 2020, McGill University for further analysis.

All microscopy, some image analysis and some cell tracking was performed at the McGill University Advanced BioImaging Facility (ABIF) *RRID* : *SCR* 017697.

