## Supplemental Table 1, Supplemental Figure 1 for "Polarity and mixed-mode oscillations may underlie different patterns of cellular migration"

### Supplementary information

| Parameter | Description | Value | Unit | References |
|---|---|---|---|---|
| $I_R$ | Basal Rac activation Rate | 0.0035 | $s^{-1}$ | [1] |
| $\delta_R$ | Rac inactivation rate | 0.025 | $s^{-1}$ | [1] |
| $L_\rho$ | Rho-dependent half-maximum inhibition of Rac | 0.34 | unitless | [1] |
| $I_\rho$ | Basal Rho activation Rate | 0.016 | $s^{-1}$ | [1] |
| $\delta_\rho$ | Rho inactivation rate | 0.016 | $s^{-1}$ | [1] |
| $L_R$ | Rac-dependent half-maximum inhibition of Rho | 0.34 | unitless | [1] |
| $\gamma$ | Ratio of total PAK to total Rac | 0.3 | unitless | [1] |
| $\delta_P$ | Paxillin dephosphorylation rate | 0.00041 | $s^{-1}$ | [1] |
| n | Hill coefficient | 4 | unitless | [1] |
| $\alpha_P$ | Linearization coefficient in paxillin activation | 2.7 | unitless | [1] |
| $L_K$ | Active PAK-dependent half-maximum activation of paxillin | 5.77 | unitless | [1] |
| $I_K$ | Additional Rac activation due to paxillin | 0.009 | $s^{-1}$ | [1] |
| $k_G$ | Association constant for GIT-PIX binding | 5.71 | $s^{-1}$ | [1] |
| $[GIT]$ | Concentration of GIT | 0.11 | $\mu M$ | [1] |
| $k_X$ | Association constant for PIX-PAK binding | 41.7 | $s^{-1}$ | [1] |
| $[PIX]$ | Concentration of PIX | 0.069 | $\mu M$ | [1] |
| $k_C$ | Association constant for $Pax_p$-GIT binding | 5 | $s^{-1}$ | [1] |
| $[Pax_{tot}]$ | Total concentration of paxillin | 2.3 | $\mu M$ | [1] |
| $\alpha_R$ | Affinity constant for PAK-RacGTP binding | 15 | unitless | [1] |
| $\epsilon$ | Time constant for $B$ | 0.01 | $s^{-1}$ | Estimated |
| $B_r$ | Resting state of $B$ | 10 | unitless | Estimated |
| $k_B$ | Recovery rate of $B$ back to its resting state | 0.04 | unitless | Estimated |
| $\gamma_R$ | Strength of $R$ feedback onto $B$ | 8.6956 | $s^{-1}$ | Estimated |
| $\eta$, $\epsilon_B$ | Parameters that guarantee the positivity of $B$ | $10^4$, $10^{-4}$ | unitless | Estimated |
| $\epsilon_L$ | Time constant of $k_B$ | $10^{-5}$ | $s^{-1}$ | Estimated |
| $\gamma_K$ | Source term | 0.15 | unitless | Estimated |

Figure S1: Summary of parameter estimations.

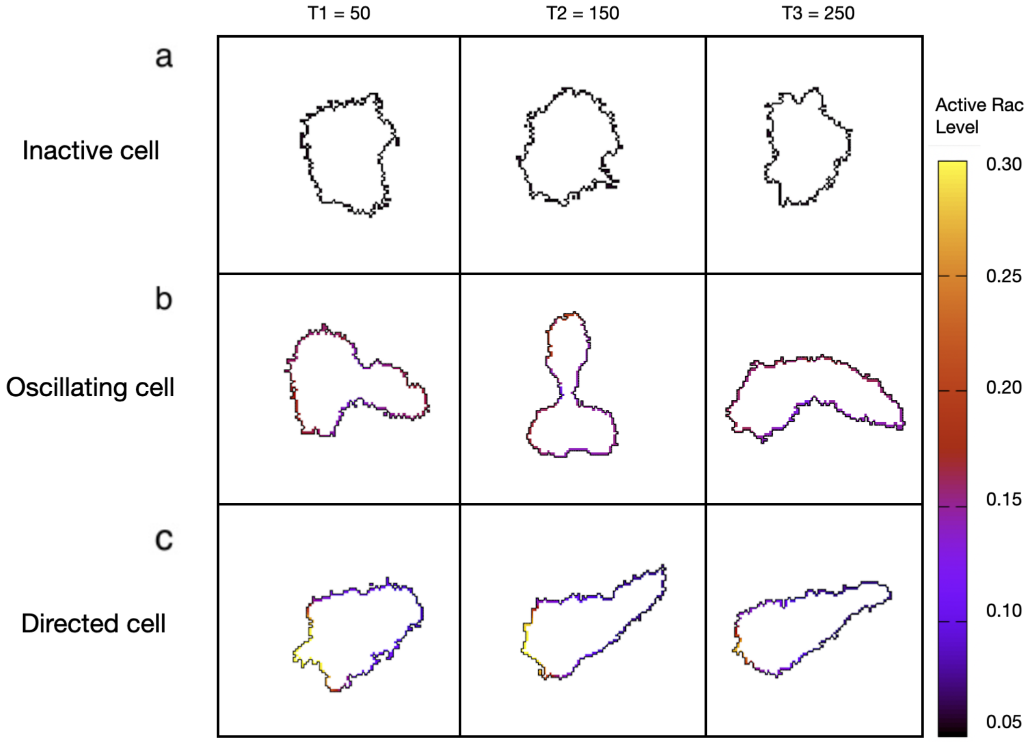


Figure S2: Simulated cells's membrane shapes for the inactive cells (a), oscillating cells (b) and directed cells (c), at three different timepoints of the simulations.
